# Backscattering amplitude in Ultrasound Localization Microscopy

**DOI:** 10.1101/2023.04.27.538586

**Authors:** Noemi Renaudin, Sophie Pezet, Nathalie Ialy-Radio, Charlie Demene, Mickael Tanter

## Abstract

In the last decade, Ultrafast Ultrasound Localisation Microscopy has taken non-invasive deep vascular imaging down to the microscopic level. By imaging diluted suspensions of circulating microbubbles in the blood stream at kHz framerate and localising the center of their individual point spread function with a sub-resolution precision, it enabled to break the unvanquished trade-off between depth of imaging and resolution by microscopically mapping the microbubbles flux and velocities deep into tissue. However, ULM also suffers limitations. Many small vessels are not visible in the ULM images due to the noise level in areas dimly explored by the microbubbles. Moreover, as the vast majority of studies are performed using 2D imaging, quantification is limited to in-plane velocity or flux measurements which hinders the accurate velocity determination and quantification. Here we show that the backscattering amplitude of each individual microbubble can also be exploited to produce backscattering images of the vascularization with a higher sensitivity compared to conventional ULM images. By providing valuable information about the relative distance of the microbubble to the 2D imaging plane in the out-of-plane direction, backscattering ULM images introduces a physically relevant 3D rendering perception in the vascular maps. It also retrieves the missing information about the out-of-plane motion of microbubbles and provides a way to improve 3D flow and velocity quantification using 2D ULM. These results pave the way to improved visualization and quantification for 2D and 3D ULM.

## Introduction

Ultrasound Localization Microscopy (ULM) recently reached unprecedented spatial and temporal resolutions in biomedical vascular imaging. Although this concept was introduced only a decade ago^1–5^, research works on ULM have been growing exponentially in the last years. The seminal paper^6^ providing proof of concept of microscopic vascular imaging at the whole organ scale introduced the key concepts used today in the field: use of ultrafast ultrasound imaging and singular value decomposition for efficiency of the microbubble signal filtering and tracking of the microbubbles, building of in-depth microscopic vascular map based on the counting of microbubbles, quantification of the bubble velocities to approximate blood flow speed and determination of the limits of resolution of the technique. This proof of concept was performed on the rat brain and applied by several research groups on various organs and animal models, such as rat kidney^7^, rabbit liver^8^, tumor models^8,9^, rat spinal cord^10,11^ and was even demonstrated in humans for the brain^12^, breast cancer^13^, liver and pancreatic tumor imaging^14^. Very recently, the technique was taken to another level with the achievement of functional ultrasound localisation microscopy, shining light on the brain neurovascular activity at a microscopic resolution^15^. To date, the vast majority of these studies have been done using 2D imaging, as both the linear transducer arrays and driving electronics are far more common and less expensive than matrix arrays, raw-column arrays, and thousand-channels or multiplexed electronics required to perform 3D ultrasound localisation microscopy^16,17^. Of course, the relative simpler implementation of 2D ultrasound localisation is at a cost, as the lack of information in the elevation direction makes velocimetry inaccurate.

In this work, we show that 2D raw ULM data gather information about the elevation direction and out-of-plane motion. This information is carried by the amplitude of the individual microbubbles backscattered signal, a parameter which was discarded so far. 2D ULM traditionally assumes that the microbubble signature on an ultrasound image resembles a local 2D point spread function (PSF), and that by retrieving the punctual 2D (x,z) center position that generated this local point spread function one can achieve imaging at a resolution below the diffraction limit. Therefore, the local microbubble signature amplitude in the (x,z) plane is used to precisely locate the (x,z) position of that microbubble. This formulation completely conceals what is an ill-posed 3D problem, with a 2D PSF (we obtain 2D images) function of a 3D space (the objects are distributed in a 3D space). This description has been already used in a very different context for ultrafast Doppler Tomography^18^. In first approximation, the (x,z) position of the microbubble signature on the image is a function of the (x,z) position of the microbubble in the 3D space, and the amplitude of the microbubble signature is a function of the (y,z) position of the microbubble in the 3D space. We show here that by considering the amplitude of the backscattered signal, we can not only obtain a different contrast more sensitive to microvessels than the usual microbubble count, but also by taking this hypothesis into account we can recover a 3D information and make velocimetry accurate.

## Results

### ULM backscattering imaging reveals a new contrast for 2D ULM and improves sensibility to small vessels

Isolated gas microbubbles used in ULM are on average Ø = 2.5 μm in diameter and relatively monodisperse (90% <8 μm)^19^, and are therefore considered as single strong Rayleigh scatterers when operating at a typical frequency of 15 MHz (Ø<λ/10). In first approximation, if we consider that the propagation occurs in a linear regime in a homogeneous medium and that all the microbubbles have the same diameter and therefore the same cross section (Sonovue cross section is 20μm^2^ for Ø = 2.2 μm, 85μm^2^ for Ø = 5 μm, therefore a ratio ∼2 in backscaterred amplitude), the backscattered energy from individual microbubbles depends only on the amplitude of the pressure wave that reaches them. Therefore, the amplitude of the signal recorded coming from a bubble is determined mainly by the propagation medium, the transducer diffraction pattern characteristics and the exact position of this microbubble within the transmitted field.

In ULM, images are conventionally obtained by counting the number of microbubbles detected in each pixel, resulting in MB count maps (often called microbubbles density maps in the literature). To image the amplitude of the backscattered signal of microbubbles in 2D ULM, we rather attribute to each pixel the average of the backscattered amplitudes (measured as the maximum of the PSF on the B-Mode image) of all the successive MBs detected in this pixel. The two resulting images are shown in figure 1.

**Figure 1:**
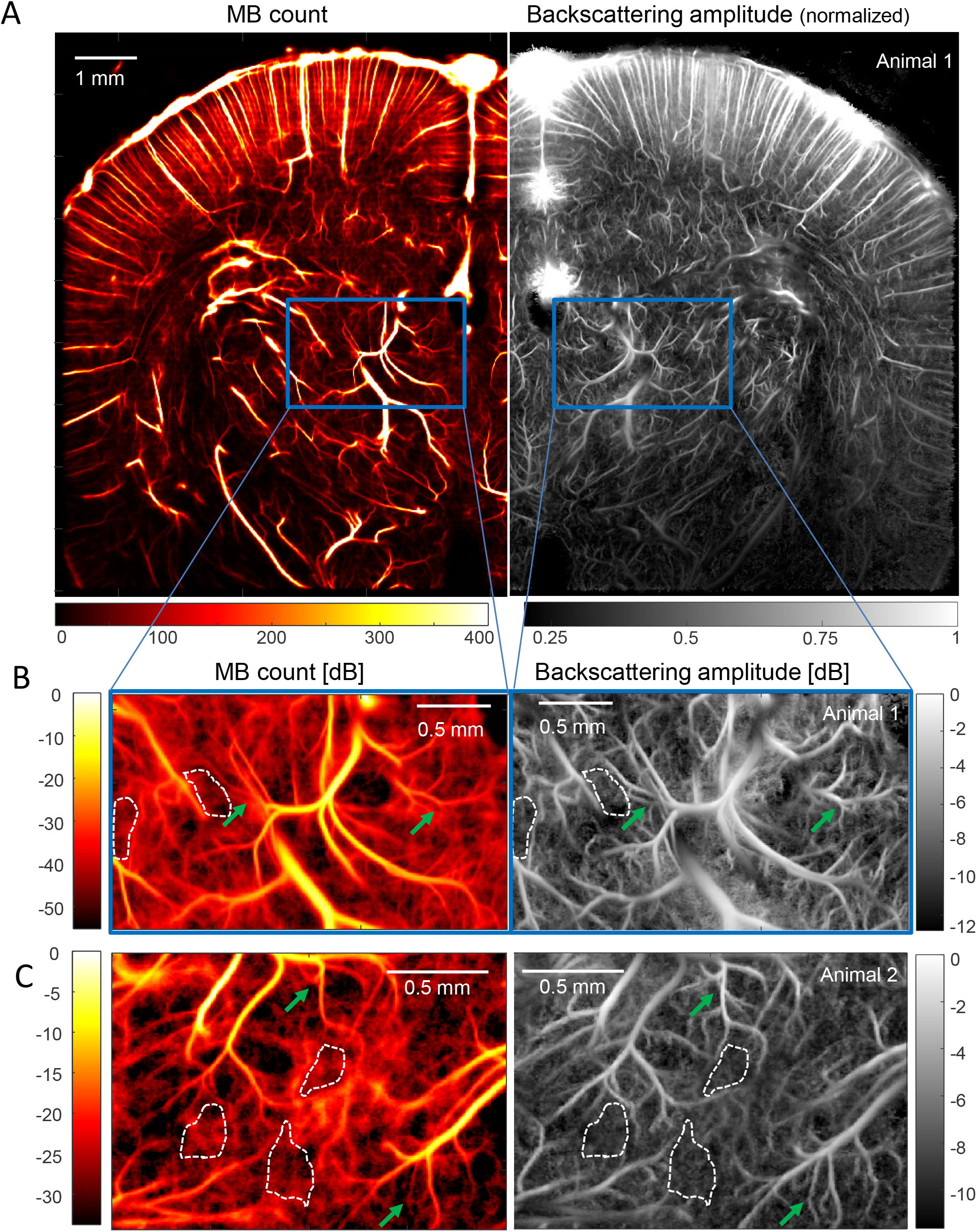
ULM Backscattering imaging offers better vessel delineation than MB count ULM. **(A)** Left: MB count ULM map of blood vessels in a rat brain coronal plane. Right: Corresponding normalized backscattering amplitude ULM map. **(C)** Zoom in the thalamus corresponding to the blue box in (A): MB count (left) and backscattering (right) maps in dB. The image log-dynamic is set by the noise level determined in areas surrounded by a white dash line. (C) Same analysis as in (B) for another thalamus area in a second animal. Green arrows depicts better small vessel delineation in backscattering imaging.

The first thing to notice in figure 1A is the difference in information content between images of MB count and backscattering amplitude. A zoom in a thalamic region of interest (blue rectangle) highlights that backscattering imaging enables a better detection of small blood vessels, as shown by the green arrows in figure 1B-C. It is worth noticing that the same detected MBs are used for the two images, it is only the mapped parameter that is different. The microbubble count and backscattering amplitude images are shown in dB with a lower bound set at their respective noise floor. Backscattering amplitude images depict well delineated vessels invisible in MB count mapping (Fig. 1B-C, green arrows).

### Backscattering from MB measured during 2D ULM imaging brings 3D information on blood vessels position in the out-of-plane direction

As the backscattered energy from individual microbubbles depends only on the amplitude of the pressure wave that reaches them, the backscattered amplitude is directly linked to the position of each microbubble within the transmit pressure field. Thus, backscattering imaging can be considered as the projection of the pressure field on the blood vessels positions. The pressure field created by the transducer (a linear array with a geometrical focusing in elevation) is represented in figure 2A (detailed simulation characterisation can be found in Fig. suppl. 1): the acoustic intensity decreases with depth (Z) because of ultrasonic attenuation along the travel path; along the out-of-plane dimension, the US beam has a certain width which varies across depth (depending on the elevation focalization and the attenuation properties). This depth dependence of the image thickness in elevation explains that more vessels are often detected in the cortical regions (before the elevation focus) compared to deeper regions (around the elevation focus) (Fig 2A), because the image thickness is larger in the near field compared to at focus. For a fixed (x_0_, z_0_) position in the imaging plane, the pressure intensity is maximal in the middle and decreases from either side (Fig 2A bottom). As a result, for a given pixel (x_0_,z_0_) in the 2D ULM image, the energy backscattered by the micro bubbles detected in that pixel (Fig 2B) depends on their position along Y within the US beam (Fig 2C).

**Figure 2:**
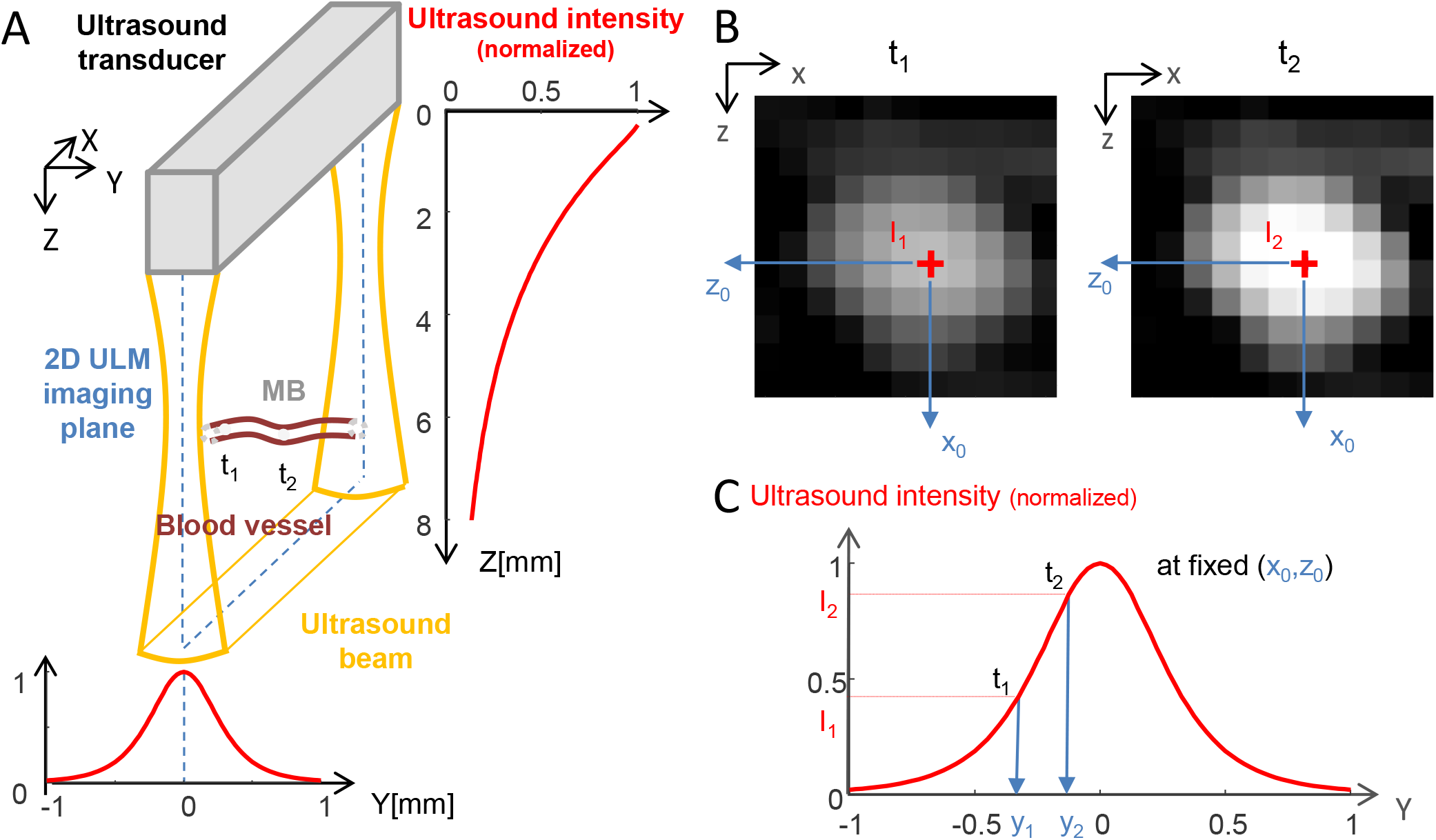
MB backscattering and ultrasound beam geometry. **(A)** Schematic of an ultrasound beam produced by a linear array transducer and of a microbubble (MB) flowing through a blood vessel orthogonal to the imaging plane. t_i_ : time point i. The red curves show the simulated normalized ultrasound intensity along the Z dimension (averaged along X and Y) (right) and y dimension (z = 5mm, averaged along X) (bottom). **(B)** BMode images of the microbubble at t_1_ and t_2_. **(C)** Variation of the backscattered ultrasound intensity and consequently of the amplitude on the BMode image as a function of Y for a fixed (x_0_,z_0_) position. The red curve show the simulated normalized ultrasound intensity along the Y dimension (z = 5mm, averaged along X).

Illustrations of this physical phenomenon are given in figure 3. The backscattering amplitude profile of a MB flowing through a blood vessel crossing the imaging plane (represented by the red line on the backscattering image in figure 3A) is given in figure 3B: as the MB enters the US beam, the incident acoustic amplitude it receives and therefore backscatters is minimal, then reaches a peak at the centre of the beam and finally decreases back as it leaves the US beam. On the backscattering images, such vessels crossing the imaging plane will appear the brightest where the vessel is in the centre of the field (green arrows), gradually darkening as the blood vessel becomes closer to the US beam edges (blue arrows). In the cortex, when imaging a coronal plane, blood vessels are almost parallel to the imaging plane so the backscattering amplitude does not vary much within the vessel. However, different vessels located at different Y positions (green, red and blue arrows on Fig. 3C) will appear in different shades of grey (whitest for the vessel (green arrow) closest to the middle of the beam, darker for the vessels closer to the beam edges (blue and red arrows). This is confirmed by the histograms of the backscattering amplitudes of the MBs travelling in those 3 vessels (Fig. 3D): MBs running through the green arrow vessel have on average higher backscattering amplitudes (green histogram). A last example is the case of blood vessels which are superimposed on a 2D ULM image, such as in figure 3E where a couple of close vessels has been highlighted with a white bar cross section. Those two vessels are slightly apart close to the cortical surface, before being completely superimposed in depth. Gathering the backscattering amplitudes of all the MBs flowing through the white segment (in Fig 3E) in a histogram reveal an obvious two-component mixture distribution of the MB backscattering amplitude (Fig. 3F), and therefore that two populations of MBs are probably flowing at different Y positions. As the orientation of arterioles and venules is well known in the cortex, it is possible to separate the MB arterial tracks from the MB venous tracks based on their axial velocity, respectively positive (downward, red) and negative (upward, blue) (Fig. 3E), and reveal that our two vessels are an arteriole and a venule.. When separating the MBs based on their arteriolar or venous nature, the two underlying backscatter amplitude distributions can be found (Fig. 3G): the venule is located closer to the centre than the artery, as the MBs from the venule display higher backscattering amplitudes.

**Figure 3:**
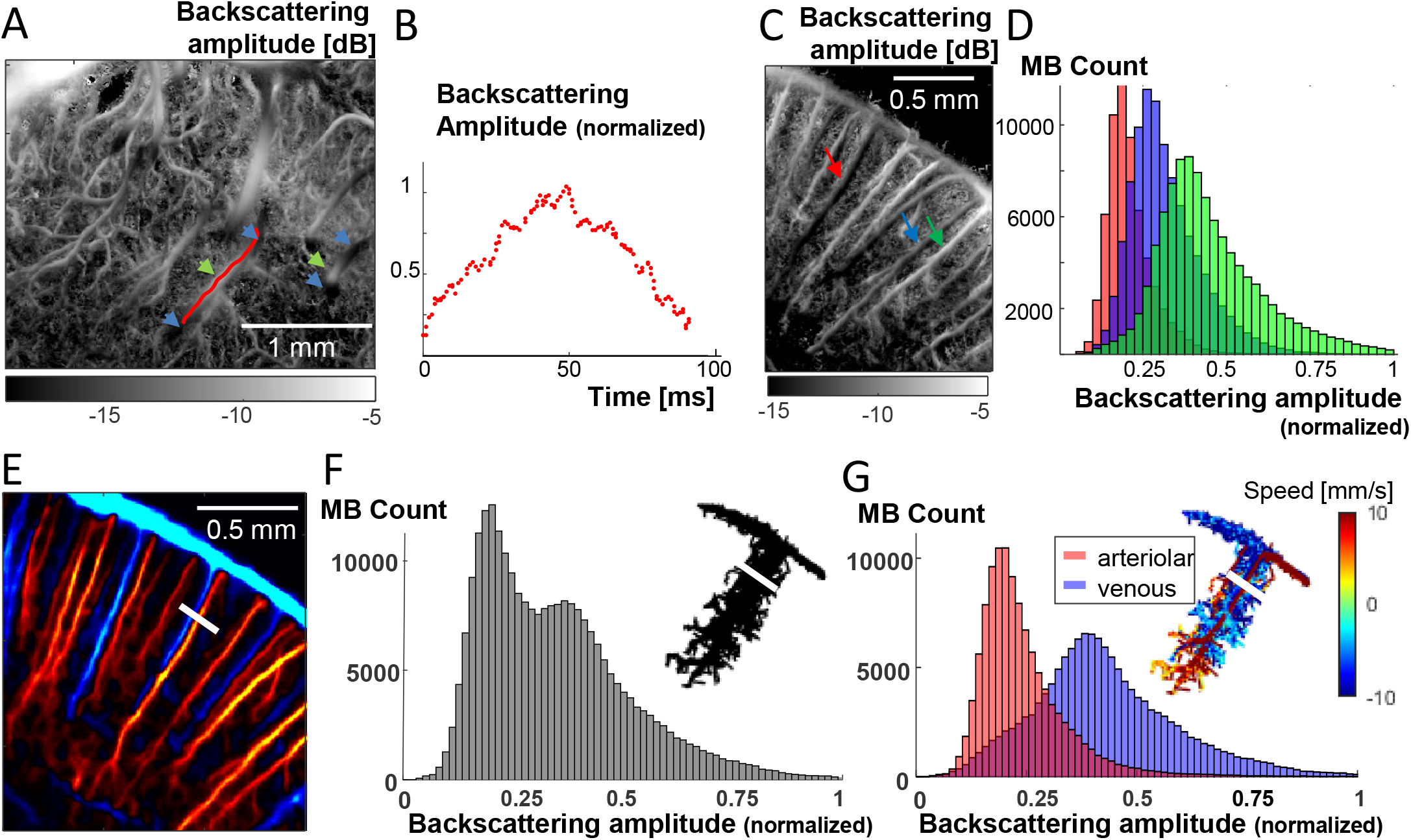
MB backscattering offers 3D information – illustrations. **(A)** Backscattering imaging ULM map of vessels in the rat thalamus crossing the ultrasound beam in the out of plane direction. The backscattering amplitude is low where the vessels are near the beam edges (blue arrows) and maximal when it is at the centre of the beam (green arrows). **(B)** Normalized backscattering amplitude as a function of time of one MB travelling in the blood vessel whose trajectory is shown on (A) as a red line. **(C)** Backscattering imaging ULM map of blood vessels in the rat cortex. **(D)** Histograms of the backscattering amplitude of all MB trajectories detected in three blood vessels pointed at by the blue, green and red arrows on (C). **(E)** ULM MB count map of a zoom in a rat cortex with downward flow in red and upward flow in blue (respectively arteriolar and venous flow). **(F)** Histogram of the backscattering amplitude of all the MB flowing through the white segment shown on (E), and related MB trajectories (top right). **(G)** Venous and arteriolar MB from (F) are separated based on flow direction and the histograms from their backscattering amplitude are displayed in blue and red respectively. Related MB trajectories with flow speed are displayed (top right).

### Backscattering imaging using 2D ULM in different planes enables 3D super-resolved localization

We then hypothesized that localizing the blood vessels in the out-of-plane direction was possible using the backscattering information available in 2D ULM imaging. To test this hypothesis, backscattering amplitude imaging of five parallel coronal planes distant from 100μm each were used (Fig. 4A). For each (x,z) pixel, a Gaussian function (see methods) is fitted on the backscattering amplitudes measured for the 5 acquisition planes (Fig. 4B). This fit is used to determine the sub-resolution position of the maximum of the backscattering amplitude function, corresponding to the position y_0_ of the blood vessel. The validity of this gaussian fitting could be confirmed by two aspects. First, the parameter w(x,z) quantifying the width of the gaussian fit can be averaged on pixels at the same depth to give w_exp_(z), which is found to be in good agreement with the ultrasound beamwidth w_sim_(z) obtained using ultrasound pressure field simulations^20,21^ with this exact ultrasound transducer (Fig. 4C). Second, the y_0_ parameter depicting the out-of-plane position of the vessel varies smoothly in vessels crossing the imaging plane, as shown on thalamic vessels in Figure 4D. This smoothness can be further increased (and i.e. the noise on y_0_ decreased) by re-estimating the positions y_0_ of the blood vessels while constraining the width of the gaussian fit to the value found experimentally w_exp_(z) (Fig. 4E) or by simulation w_sim_(z) (Fig. 4F), with a good agreement between the two methods.

**Figure 4:**
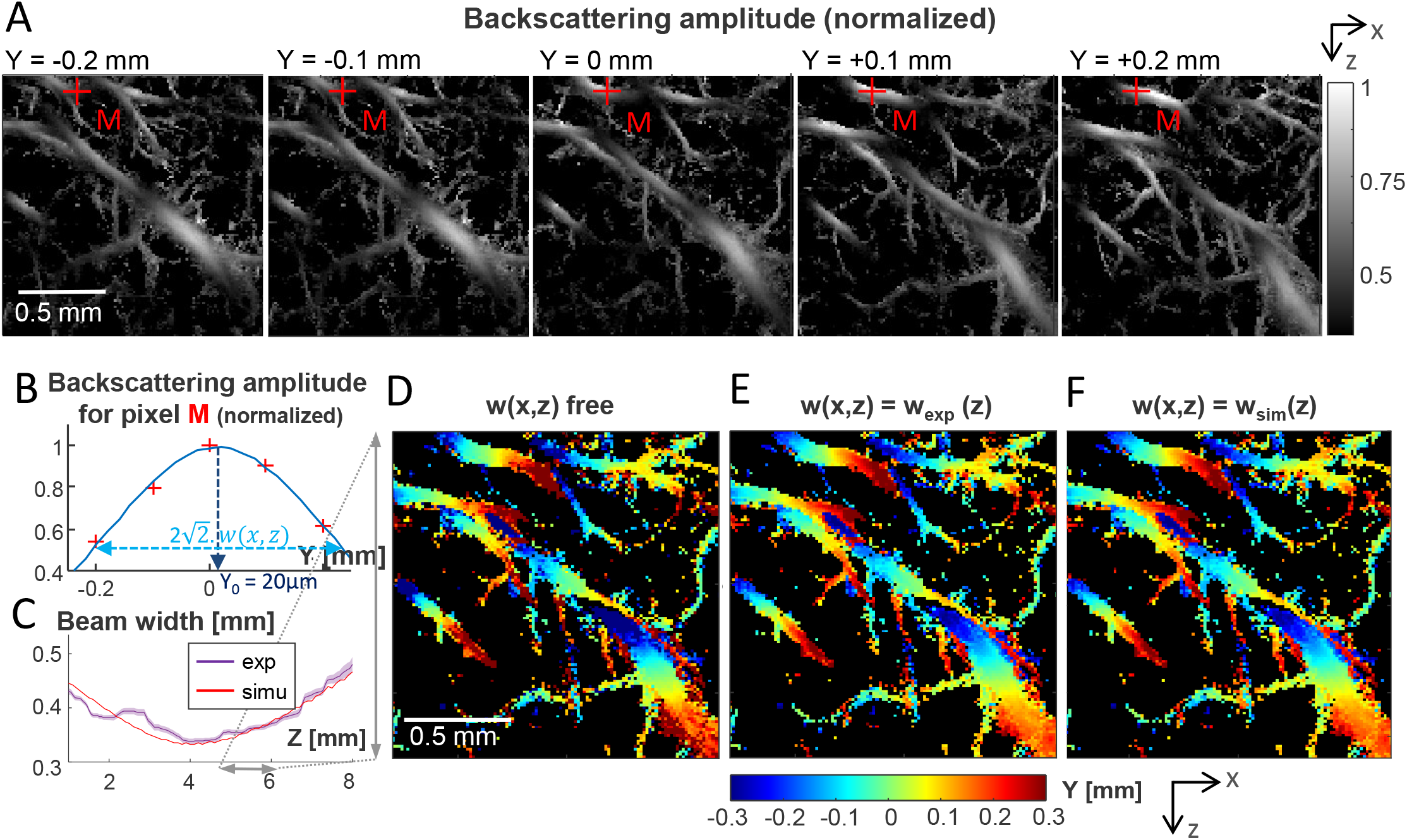
2D ULM backscattering imaging and 3D sub-resolution localization. **(A)** 2D ULM backscattering imaging of 5 coronal planes in the rat thalamus spaced by 100μm from each other in the elevation (Y) direction. **(B)** Backscattering amplitude of pixel M (red cross in (A)) for the 5 planes. The blue curves is a gaussian fit on the 5 points. y_0_ depicts the sub-resolution localisation of the maximum, i.e. the y-position of the vessel. **(C)** Violet curve: experimental gaussian fitting width parameter averaged per depth (mean+/-SEM). Red curve : beam width obtained by simulation. **(D-F)** Sub-resolution maps of the location (y_0_) in the Y (out-of-plane) direction based on a gaussian fit (width parameter = w). (D) No constraint on w. (E) Constrained width parameter w(z) obtained from experiment results (violet curve in C). (F) Constrained width parameter w(z) obtained from transducer beam simulation (red curve in C).

### Backscattering imaging in 2D ULM retrieves 3D speed vector estimations

Another source of bias in 2D ULM is that the blood velocity can only be estimated from the two in-plane components of the speed vector. The missing out-of-plane component of the speed vector is responsible for the underestimation of the true speed, especially for blood vessels crossing the imaging plane. Here, we use the backscattering information of a **single plane** acquisition to access the out-of-plane component (Fig. 5A-D) using a simplified framework. The in plane 2D component of the flow speed is determined classically using the differential of the successive MB positions in time (Fig. 5A), while the out of plane component is deduced from a modelling of the out-of-plane angle of the vessel: assuming a rectilinear crossing of the US beam by the vessel (as depicted by the maroon line in Fig. 5B), we can estimate the angle between this blood vessel and its projection on the imaging plane from the backscattering profile along this vessel (Fig 5C) and the knowledge of the beam width. We did it for 4 distinct vessels using a single 2D ULM acquisition corresponding to the central plane of the 5-plane experiment. We obtained a good agreement between the angles found from the single plane ULM acquisition and the angles obtained using the multiplane experiment (Fig. 5D).

**Figure 5:**
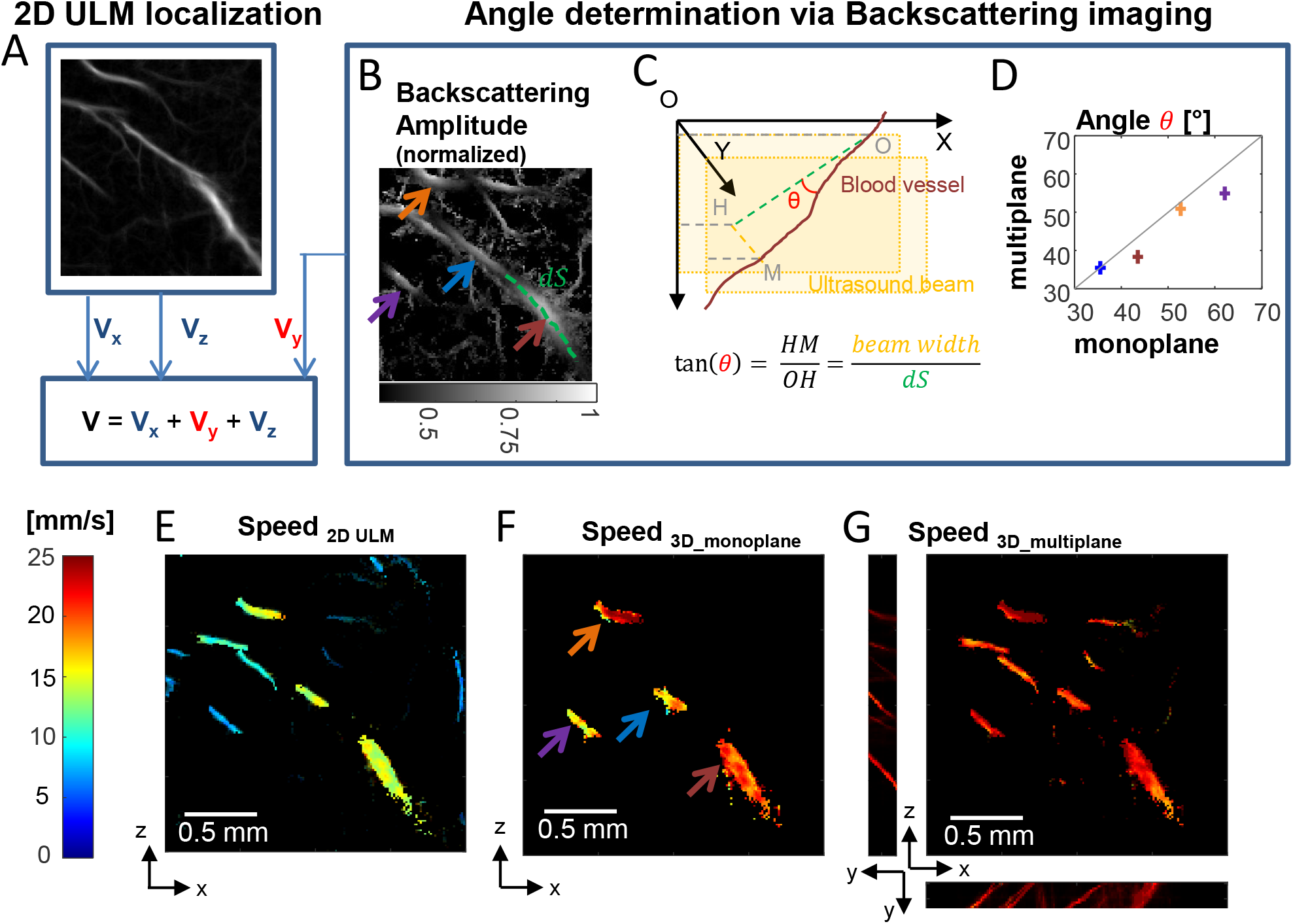
Backscattering imaging and speed vector correction. **(A-D)** Schematic of the speed vector correction : **(A)** V_x_ and V_y_ are estimated thanks to 2D ULM while Vy is estimated thanks to ULM backscattering imaging **(B)**. Arrows depict selected vessels for (D). **(C)** Blood vessel angle *θ* toward imaging plane is determined using knowledge of the local beam width and width at half maximum dS of backscattering amplitude function along the vessel. The schematic explains the geometry. **(D)** Comparison between angle estimation using one plane and multiple planes backscattering information. **(E)** Speed estimation in classical 2D ULM. **(F)** Speed estimation correction with ULM backscattering imaging using a single plane. **(G)** For comparison, speed estimation correction using ULM backscattering imaging information in multiple planes, as done in figure 4.

The speed vector can then be corrected thanks to the knowledge of this angle. A comparison between the estimation of speed usually done in 2D versus the corrected speed using the backscattering profile based on a single plane acquisition is given in figure 5E-F. We also show as a ground truth the speed estimation using the 3D localization thanks to the 5-plane experiment (Fig. 5G). The velocity was largely underestimated when using 2D ULM only (Fig. 5E) compared to when corrected using the backscattering amplitude information in one plane only (Fig.5f). We founnd that for the four vessels investigated, the error on the speed was 100+/-80% using conventional 2D ULM, but reduced to 20+/-20% after correction using the monoplane speed correction.

## Discussion

Backscattering imaging, a contrast overlooked until now, provides a new informative parameter in Ultrasound Localization Microscopy. First, backscattering imaging delineates more small vessels than conventional ULM images (Fig. 1B-C) due to an intrinsic lower spatial noise. But these images also present an interesting 3D rendering effect as the backscattering amplitude intrinsically conveys physical information about the out-of-plane distance. Finally, and even more importantly, this backscattering amplitude can be exploited to recover the missing information about motion in the out-of-plane direction (Fig. 5), even with a single plane 2D ULM acquisition.

Backscattering ULM imaging is less sensitive to noise and better detects those small vessels compared to conventional MB count ULM. In MB Count maps, the signal is driven by blood flow (the bigger the blood flow, the more MBs are detected during the acquisition time). In backscattering imaging, the signal is mainly driven by the position of the blood vessels within the ultrasound beam independently from hemodynamics. One should notice that although these images are extracted from the same raw data, the noise signal due either to false or missed detections has a lower influence in backscattering imaging than in conventional ULM: respectively they will be averaged out in the estimation of the backscattering amplitude signal, while they will extensively add noise to the value of the MB count. Backscatter imaging could also be more adapted for segmentation algorithms and could provide better insights on the size and order of branching capillaries detected using ULM imaging.

Some assumptions were made in this work. We considered that all the MBs are identical scatterers relying on similar backscattering properties, which is surely not entirely the case. However, the images are made by averaging the backscattering amplitudes from all the MBs randomly detected in the pixel, which will smooth out this effect. Similarly to 2D ULM localization, the hypothesis that the individual MBs are completely isolated might not always be valid, particularly for large vessels. It is even more likely along Y elevation direction as the diffraction limit set by the US beam focalization is around 500μm in the elevation direction in comparison to 100μm within the imaging plane. This might be the reason why backscattering amplitude rockets in very large vessels (Fig. 1).

Of course, 3D ULM imaging can be achieved using either fully populated 2D matrix array technologies^17^ or Raw Column Arrays (RCA) technologies^22–25^. However, to date, conventional linear arrays are mainly used for ULM imaging because they are cheaper, simpler and have a much better sensitivity compared to matrix arrays. The backscattering ULM imaging approach proposed here could deliver more accurate speed quantifications for those 2D acquisitions. For multi plane experiments, the localization technique combined with backscatter amplitude imaging provides 3D volumetric images with a sub-wavelength resolution in the out-of-plane direction. Using one single plane acquisition, the symmetry of the problem along the elevation direction does not permit to retrieve the exact location of the microbubble in the out-of-plane direction (below or above the imaging plane) but only its absolute distance to the imaging plane. Nevertheless, the 3D correction of the speed vector does not require this knowledge and although it was presented in a very simple situation, it could be further refined to offer reliable speed quantification for 2D-ULM.

We mainly investigated the 3D / US beam influence on the backscattering ULM images. However, other differences potentially affecting the backscatter amplitude within the vascular network for instance in pressure (veinous vs arteries) could also potentially influence the backscattering properties and could be further investigated.

## Methods

### Animals

All experiments were performed in compliance with the European Community Council Directive of September 22, 2010 (010/63/UE) and were approved by the french Ethics Committee (Comité d’éthique en matière d’expérimentation animale N°59, ‘Paris Centre et Sud’, project #2017-23). All methods are reported in accordance with ARRIVE guidelines. Accordingly, the number of animals in our study was kept to the necessary minimum following the 3Rs (Reduce, Refine, Replace) guidelines. Experiments were performed on N=3 male (Age 7-9 weeks) Sprague-Dawley rats (Janvier Labs; Le Genest St Isle, France), weighing 200-300g at the beginning of the experiments. Animals (housed two per cage) arrived in the laboratory one week before the beginning of the experiment and were maintained under controlled conditions (22 ± 1°C, 60 ± 10% relative humidity, 12/12h light/dark cycle, food and water ad libitum). At the end of the experiments, animals were sacrificed under anaesthesia with a quick cervical dislocation. All animals included in this study were untreated and were used randomly in the various experiments.

### Surgical procedure and preparation for imaging

Under deep anaesthesia (intraperitoneal (IP) bolus of medetomidine (Domitor, 0.4 mg.kg) and ketamine (Imalgène, 40 mg.kg^-1^)), a catheter filled with saline was inserted in the jugular vein of the rat before positioning the animal on the stereotaxic frame. A craniotomy (removal of the skull) was then performed between Bregma and Lambda. During the surgical procedure and the imaging session, the animals’ body temperature was kept at 37°C using a heating blanket and an intra-rectal probe (Physitemp, USA). Around 45min after induction (when the craniotomy was completed), the anaesthesia was maintained but reduced, using subcutaneous perfusion of Medetomidine (0.1 mg/kg/h) and ketamine (12.5 mg/kg/h) with a syringe pump. The heart and respiratory frequencies were monitored continuously to ensure stability of the anaesthesia (Labchart, AD Instruments). Each imaging session lasted between two and four hours. Two millilitres of saline were gently dropped on to the brain (the dura mater was kept intact), followed by echographic gel (Dexco Medical, France). The ultrasonic probe was then positioned just above the window, using a three-axis motorized system on which the ultrasound probe was fixed.

### Ultrasound data acquisisiton

ULM acquisitions were performed similarly to the methods described by Demené et al., 2021^12^, but using a continuous injection at a 3.5mL/h dose of Sonovue (Braco, Italy, reconstructed in 5mL of saline, as recommended by the manufacturer), using a push syringe (KD Scientific, USA). A magnet was inserted in the syringe in order to mix the microbubbles solution during the acquisition. Continuous injections enabled a stable number of MBs to be secured for more than 20 minutes with approximatively 30 MB per ultrasound frame after detection and tracking. Blocks of 400 compounded frames at a 1000Hz framerate (each frame is a compound image acquired with angles at -5°, -2°, 0°, +2°, +5° fired at a 5000Hz PRF) were acquired using the same probe (128 elements, 15 MHz, 110 μm pitch, 8 mm elevation focus, Vermon, Tours, France). The pulse shape corresponds to 2 periods of sinusoids, the transmit voltage is 5V, the Mechanical index is 0.09.

For the multiplane experiments, the probe was moved along the rostro-caudal axis with a 100μm step between each of the 5 planes. A block of 400 compounded frames was acquired for the first bloc before moving to the second block, and so on, with a 150ms pause to let enough time for the motors to move. At the end of the 5^th^ plane the probe was moved back to the first plane. This was repeated so that 525 blocks of 400ms were acquired for each plane, leading to a 210 seconds accumulation time for the ULM images.

### Ultrasound Localization Microscopy processing

Beamformed data were filtered using the SVD spatio-temporal clutter filter^26^ to discriminate the ultrasonic signature of individual microbubbles from tissue signals; the 10 first singular values were discarded. Images were interpolated (Lanczos interpolation kernel) down to (probe spatial picth/6 x λ/6). A binary mask was built based on the vesselness filtering^27^ of this stack of images (3D implementation available on Mathworks file exchange, ©Dirk-Jan Kroon2009, and ©Tim Jerman, 2017). Microbubbles were detected as the brightest local maxima with high correlation (> 0.5) with a typical point spread function (imaging response of an isolated microbubble, modelled as a Gaussian spot of axial and lateral dimension of lambda). Sub-pixel maxima localization was then performed using a fast local second-order polynomial fit. The resulting coordinates were rounded to the chosen pixel size (here 13.75x12.5μm = initial pixel size/8). Tracking of the maxima positions was performed using a classical particle tracking algorithm (simpletracker.m available on Mathworks ©Jean-Yves Tinevez, wrapping matlab munkres algorithm implementation of ©Yi Cao 2009), with no gap filling and maximal linking distance corresponding to a 100mm/s maximum speed. Only tracks with MB detected in at least 10 successive ultrafast frames were selected. The successive positions gathered in one track were used to compute the interframe bubble velocity vector components (along probe x-axis and depth z-axis) and absolute velocity magnitude. We added a linear spatial interpolation on each track to count one MB detection in every pixel on the MB path. Maps of MB Count were computed by counting all the MBs detected in one pixel during the acquisition time; velocity maps were computed as their mean velocity. In plane pixel size for image reconstruction is ∼13μm (13.75x12.5μm). To take into account unavoidable motion drift occurring on time scales much slower than the cardiac or breathing time scales, we also added an intensity-based spatial registration (translation transformation) based on 10s ULM MB Count maps (dimension x,z,t) to correct for any drift during the global acquisition time (>20min). We used Matlab functions *imregconfig* (monomodal option) and *imregtfrom* (translation option).

### Backscattering imaging

During the processing to create ULM images, microbubbles were detected as brightest local maxima with a good correlation with the PSF on BMode images. The value of this maxima on the BMode images were extracted and associated with each localized MB detection. This value is referred as the backscattering amplitude. Then, instead of attributing to each ULM map’s pixel the number of MB detected in this pixel during the whole acquisition, we attribute to each pixel the mean value of the backscattering amplitude of at the MBs detected in that particular pixel. The grid chosen is the same as the one chosen for the MB Count and Velocity maps.

### MB backscattering time courses and histograms

For the backscattering time courses and histograms, shown in Fig 3 and Fig 5C, MB tracks from particular blood vessels are isolated by choosing a segment transversal to the blood vessel and selecting one or all the tracks passing through this segment. The backscattering amplitude of those MB are used to create the histograms and time courses.

### Multiplane elevation localization

For each of the 5 planes, a backscattering ULM map is created. As the planes are separated from 100μm and the beam width is over 500μm, the same blood vessels are present on the different ULM maps. For each pixel, a gaussian fit 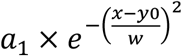 is performed on the 5 backscattering amplitudes from the 5 planes (Fig 4B). The position of the blood vessel corresponding to the pixel in the elevation direction (Y) is given by the y position where this fit is maximum: y_0_. This fit is supposed to correspond to the US amplitude field along Y for each (x,z) position. *a*_*1*_ corresponds to the amplitude and is supposed to decrease along the depth due to ultrasonic attenuation, w corresponds to the width of the beam, which is the largest near the transducer, converges towards the focal depth where it is minimal and then diverges. The resulting localisation is displayed Fig 4D.

We consider that the US intensity field is the same at each depth (which is roughly true, except close from the edges of the plane in the X direction), the beam width *w*_*exp*_ is estimated by averaging along each depth all w measured for each pixel. Outliers are first excluded. Fig 4C displays the resulting experimental width in purple (mean+-SEM). A median filter (was first applied along the Z depth direction).

For each pixel, we then perform again a fit on the 5 backscattering amplitudes, but constraining the w by the experimental measurement *w*_*exp*_ (Fig 4E). We also do it by constraining with the w value obtained by the simulation *w*_*sim*_ (linked to the Full Width Half Maximum (FWHM) of the simulated ultrasound intensity profile along Y by 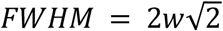), shown in Fig 4F.

### Speed correction in multiple planes acquisitions

Now that we have the 3D localization of the blood vessels, we attribute to each MB detection its y localization depending on the position of the blood vessel it belongs to at the (x,z) position it was detected. We can measure the angle between the vessels and the imaging plane knowing the y positions. We can also run again the speed calculation for every track but this time also using the y component.

### Speed correction in single plane acquisitions

We first extract the tracks which belong to the blood vessel we want to correct the speed estimation of, and then obtain the backscattering profile along the vessel’s centerline. We calculate the angle *θ* as 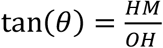 (see Fig 4C for the geometric construction). OH is evaluated by the FWHM in curvilinear abscissa of the backscattering profile along the centerline. HM, the beam width, is evaluated by the FWHM of the simulated ultrasound intensity profile. We then compute the corrected speed by: 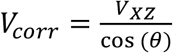 where V_XZ_ is the speed calculated in conventional 2D ULM.

### US beam simulation

The spatial ultrasound field transmitted by the ultrasonic probe is simulated using Field II software^20,21^. The simulation parameters are described in Table 1. An apodization along elevation is modelled as an Hanning window scaled between 0.4 and 1, and along *x* as a Kaiser window.

**Table 1.** Ultrasonic and simulation parameters used for the modelling of the ultrasonic beam patterns

| Ultrasound probe parameters |  | Simulation parameters |  |
| --- | --- | --- | --- |
| Frequency | 15.625 MHz | Speed of sound | 1540 m/s |
| Nb piezo elements | 128 | Attenuation | 134.24 Np/m (value for brain white matter at 15.625MHz © 2010-2022 IT'IS Foundation) |
| Dimension piezo elements: |  | Spatial sampling: |  |
| X | 108µm | X | 100µm |
| Y | 1.5mm | Y | 10µm |
|  |  | Z | 100µm |
| Inter element distance (kerf) | 2 µm | Sampling frequency | 300 MHz |
| Elevation focus | 8 mm | Input waveform | 2 periods of sinusoid |

The 3D (xyz) pressure field is simulated in space and time after emission of a plane wave with the said excitation by the defined transducer array. In each point of space (x,y,z) is therefore recorded a pressure signal p(t). In that simple case, the amplitude of this p(t) can be seen as proportional to an amplitude antenna gain. To account for antenna gain in both transmission and reception, this pressure is squared, before taking the maximum value in time. Once normalised in space, this quantity defines the simulated backscattering ultrasound intensity that is displayed in figure 2 and used to determine the simulated beam width on several occasions throughout the article.

### Data availability statement

The datasets used and analyzed during the current study available from the corresponding author on reasonable request.

### Code availability statement

Custom codes used for the collection and analysis of the data used in this study are protected by Inserm and can only be shared upon request, with the written agreement of Inserm.

## References

1. Couture, O., Besson, B., Montaldo, G., Fink, M. & Tanter, M. Microbubble ultrasound superlocalization imaging (MUSLI). in 2011 IEEE International Ultrasonics Symposium 1285–1287 (2011). doi:10.1109/ULTSYM.2011.6293576.

2. Siepmann, M., Schmitz, G., Bzyl, J., Palmowski, M. & Kiessling, F. Imaging tumor vascularity by tracing single microbubbles. in 2011 IEEE International Ultrasonics Symposium 1906–1909 (2011). doi:10.1109/ULTSYM.2011.0476.

3. Desailly, Y., Couture, O., Fink, M. & Tanter, M. Sono-activated ultrasound localization microscopy. Applied Physics Letters 103, 174107 (2013).

4. Viessmann, O. M., Eckersley, R. J., Christensen-Jeffries, K., Tang, M. X. & Dunsby, C. Acoustic super-resolution with ultrasound and microbubbles. Phys Med Biol 58, 6447–6458 (2013).

5. O’Reilly, M. A. & Hynynen, K. A super-resolution ultrasound method for brain vascular mapping. Med Phys 40, 110701 (2013).

6. Errico, C. et al. Ultrafast ultrasound localization microscopy for deep super-resolution vascular imaging. Nature 527, 499–502 (2015).

7. Foiret, J. et al. Ultrasound localization microscopy to image and assess microvasculature in a rat kidney. Scientific Reports 7, 13662 (2017).

8. Zhang, W. et al. Super-Resolution Ultrasound Localization Microscopy on a Rabbit Liver VX2 Tumor Model: An Initial Feasibility Study. Ultrasound in Medicine & Biology 47, 2416–2429 (2021).

9. Lowerison, M. R., Huang, C., Lucien, F., Chen, S. & Song, P. Ultrasound localization microscopy of renal tumor xenografts in chicken embryo is correlated to hypoxia. Sci Rep 10, 2478 (2020).

10. Sloun, R. J. G. van et al. Super-Resolution Ultrasound Localization Microscopy Through Deep Learning. IEEE Transactions on Medical Imaging 40, 829–839 (2021).

11. Claron, J. et al. Large-scale functional ultrasound imaging of the spinal cord reveals in-depth spatiotemporal responses of spinal nociceptive circuits in both normal and inflammatory states. PAIN 162, 1047–1059 (2021).

12. Demené, C. et al. Transcranial ultrafast ultrasound localization microscopy of brain vasculature in patients. Nature Biomedical Engineering 5, 219–228 (2021).

13. Opacic, T. et al. Motion model ultrasound localization microscopy for preclinical and clinical multiparametric tumor characterization. Nature Communications 9, (2018).

14. Huang, C. et al. Super-resolution ultrasound localization microscopy based on a high frame-rate clinical ultrasound scanner: an in-human feasibility study. Phys. Med. Biol. 66, 08NT01 (2021).

15. Renaudin, N. et al. Functional ultrasound localization microscopy reveals brain-wide neurovascular activity on a microscopic scale. Nat Methods 19, 1004–1012 (2022).

16. Heiles, B. et al. Ultrafast 3D Ultrasound Localization Microscopy Using a 32 $\times$ 32 Matrix Array. IEEE Transactions on Medical Imaging 38, 2005–2015 (2019).

17. Demeulenaere, O. et al. Coronary Flow Assessment Using 3-Dimensional Ultrafast Ultrasound Localization Microscopy. JACC: Cardiovascular Imaging 15, 1193–1208 (2022).

18. Demené, C. et al. 4D microvascular imaging based on ultrafast Doppler tomography. NeuroImage (2015) doi:10.1016/j.neuroimage.2015.11.014.

19. Schneider, M. Characteristics of SonoVueTM. Echocardiography 16, 743–746 (1999).

20. Jensen, J. FIELD: A program for simulating ultrasound systems. Medical and Biological Engineering and Computing 34, 351–352 (1996).

21. Jensen, J. A. & Svendsen, N. B. Calculation of pressure fields from arbitrarily shaped, apodized, and excited ultrasound transducers. IEEE Transactions on Ultrasonics, Ferroelectrics, and Frequency Control 39, 262–267 (1992).

22. Jensen, J. A. et al. Three-Dimensional Super-Resolution Imaging Using a Row–Column Array. IEEE Transactions on Ultrasonics, Ferroelectrics, and Frequency Control 67, 538–546 (2020).

23. Sauvage, J. et al. A large aperture row column addressed probe for in vivo 4D ultrafast doppler ultrasound imaging. Phys Med Biol 63, 215012 (2018).

24. Flesch, M. et al. 4D in vivo ultrafast ultrasound imaging using a row-column addressed matrix and coherently-compounded orthogonal plane waves. Phys Med Biol 62, 4571–4588 (2017).

25. Jensen, J. A. et al. Anatomic and Functional Imaging Using Row-Column Arrays. IEEE Trans Ultrason Ferroelectr Freq Control 69, 2722–2738 (2022).

26. Demené, C. et al. Spatiotemporal Clutter Filtering of Ultrafast Ultrasound Data Highly Increases Doppler and fUltrasound Sensitivity. IEEE Transactions on Medical Imaging 34, 2271–2285 (2015).

27. Jerman, T., Pernuš, F., Likar, B. & Špiclin, Ž. Enhancement of vascular structures in 3D and 2D angiographic images. IEEE Trans. Med. Imaging 35, 2107–2118 (2016).

