## Supplementary Figures for "Backscattering amplitude in Ultrasound Localization Microscopy"

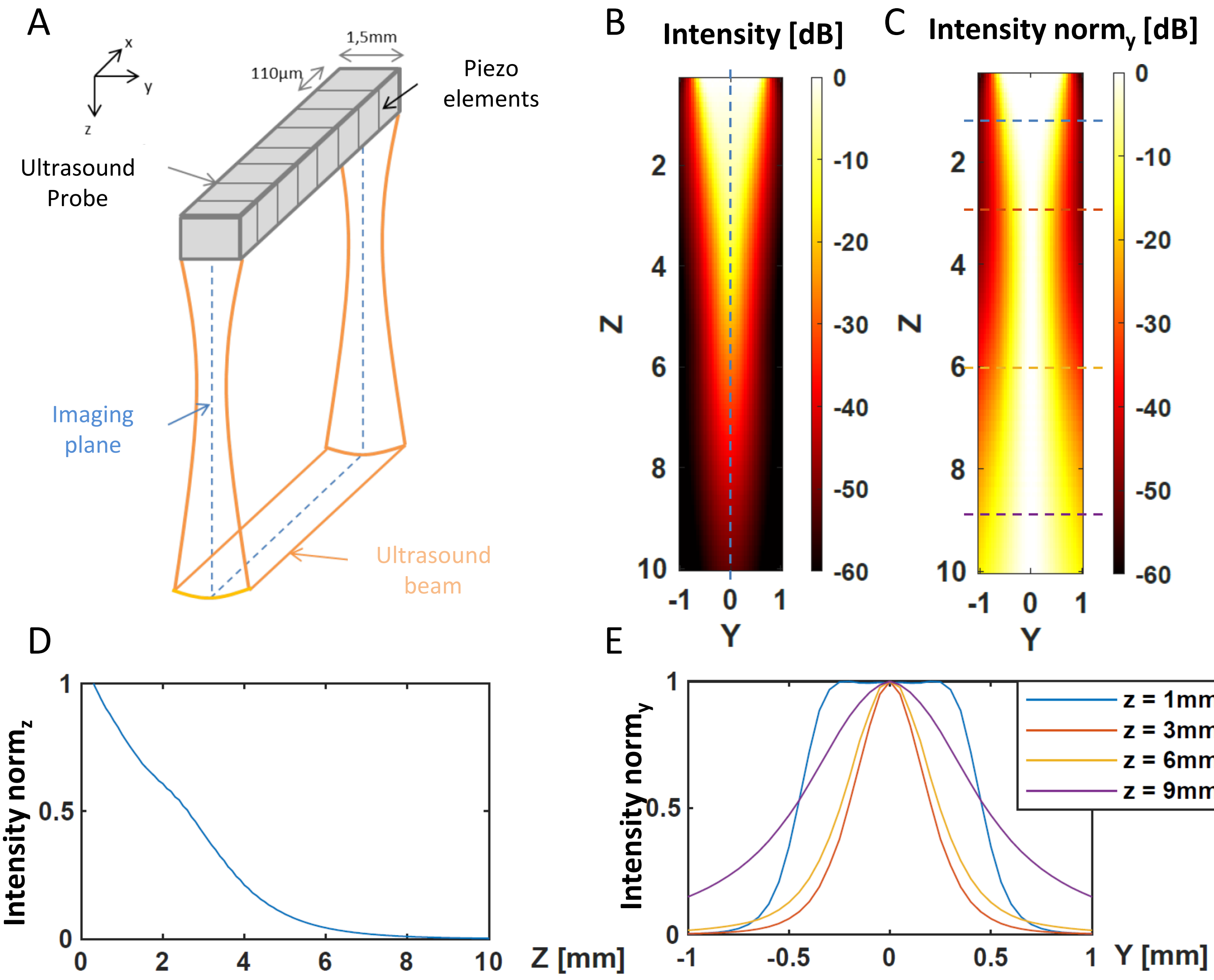


**Figure suppl 1 : Simulation of the ultrasound beam pressure field and Intensity. (A)** Schematic of the ultrasound probe and characteristic spatial dimensions. **(B)** Ultrasound Intensity in the YZ plane at the X central position (X = 0). **(C)** Ultrasound Intensity in the YZ plane (X = 0) normalized for each depth z. **(D)** Decrease of ultrasound intensity with depth z (**I**(x=0, y=0, z) along the blue dashed line in B). **(E)** Ultrasound intensity profile along y for a given depth, normalized for each depth as in C (along colored lines in C).
